# Recombinant production of rabbit β-Nerve Growth Factor and its biological effect on rabbit sperm

**DOI:** 10.1101/458612

**Authors:** Ana Sanchez-Rodriguez, Paloma Abad, Maria Arias-Alvarez, Pilar G. Rebollar, José M. Bautista, Pedro L. Lorenzo, Rosa M. Garcia-Garcia

## Abstract

The neurotrophin β-Nerve Growth Factor (β-NGF) is flourishing as a protein with important roles in the ovulation induction process in induced-ovulation species but data in rabbits are still inconclusive, probably due to the species-specificity effect of the neurotrophin to trigger the ovulation. Moreover, β-NGF seems to have a role in sperm function. To clarify these functionalities we aimed, in the present research: 1) to newly synthesize a functional recombinant β-NGF from rabbit (rrβ-NGF), 2) to reveal differences in the amino acid sequence of rabbit β-NGF compared to other sequences of induced and spontaneous ovulator species, and 3) to assess the effects of rrβ-NGF on sperm viability and motility. The nucleotide sequence of *NGF* from rabbit prostate was sequenced by Rapid Amplification of cDNA Ends (RACE) and annotated in GenBank (KX528686). Then, rrβ-NGF was produced in CHO cells and purified by affinity chromatography. Western blot and MALDI-TOF analyses confirmed the correct identity of the recombinant protein. rrβ-NGF functionality was validated in PC12 cells through a successful dose-response effect along 8 days. The comparison of the amino acid sequences of NGF between rabbit and other species suggested some relevant substitutions at its binding site to both the high-(TrkA) and the low-(p75) affinity receptors. The addition of rrβ-NGF in rabbit sperm, in a time- and dose-response study, did not affect its viability but slightly changed some of its motility parameters at the highest concentration used (100 ng/ml). Thus, it can be considered that this new recombinant protein may be used for biotechnological and reproduction assisted techniques in ovulation-induced species.

## Introduction

Beta Nerve Growth Factor (β-NGF) is a neurotrophin first described for its function in survival and maintenance of sympathetic and sensory neurons in the nervous system [1]. Similar to many other growth factors, β-NGF is synthesized as a precursor (pro-NGF), which is a 241-residue protein composed by a signal peptide of 18 residues, a pro-peptide of 103 residues and the mature form of 120 residues in the C-terminal end (data extrapolated and generalized from human β-NGF: http://www.uniprot.org/uniprot/P01138). N-glycosylation of pro-NGF is essential for the processing, secretion and construction of the tertiary fold of the homodimer of β-NGF [2, 3]. Then, the protein should form a dimer whose correct folding is crucial for the performance of the biological activity [4]. Furthermore, the post-translational modifications of the molecule, i.e. three disulfide bonds and the cysteine knot within the two β-NGF chains, are essential for the correct structure [5]. Since only intracellular engineering of mammalian cells is able to efficiently generate all these modifications, Chinese Hamster Ovary (CHO) cells have been used elsewhere for the recombinant production of human and mouse β-NGF [3, 6], resulting in biologically active proteins.

β-NGF achieves its function when it binds to its high-affinity receptor, TrkA, present in a wide range of cells, such as neurons [7], keratinocytes [8], cardiac myocytes [9] or cells of sexual organs [10, 11], developing different actions depending on where is located. This receptor is also present in the cell surface of PC12 cells, a cell line derived from pheochromocytoma of the rat adrenal medulla, extensively used for testing the biological activity of β-NGF [12]. These cells respond to β-NGF and they turned from proliferating chromaffin-like cells to non-dividing sympathetic-neuron-like cells that extend axons or neurites [13]. This effect is produced when β-NGF binds to TrkA and mediates signaling cascades, such as ERK/MAPK and PI3K/AKT pathways [14].

Recently, β-NGF has been identified as the ovulation-inducing factor (OIF) in the seminal plasma of some species of reflexive ovulation, as camelids [15–18]. In rabbits, another induced ovulatory species, the protein and its receptors (TrkA and p75) have been identified in all reproductive male and female tissues (testes, epididymis and accessory glands of male rabbits [10, 11, 19, 20], and uterus, oviduct and ovary [21, 22, 23]) as well as in seminal plasma [10, 11, 24]. However, there are no studies which demonstrate that β-NGF is the OIF in this species, although some kind of ovarian response in terms of the increase of hemorrhagic follicles (non-ruptured anovulatory follicles full of blood) have been evidenced using β-NGF of murine species [11], or rabbit SP [16, 25] intramuscularly injected. Although β-NGF is a highly conserved protein among species [26], a detailed comparison analysis of the amino acid sequences between induced- and spontaneous-ovulation species, could help to identify different ovulation patterns among species.

In addition, β-NGF has been detected in semen in some other species but its role in sperm physiology is barely studied. This neurotrophin has been localized in round spermatids and spermatocytes in the reproductive tract in mouse and rat [27, 28] and it is suggested to intervene in sperm maturation [29, 30]. In ejaculated sperm, β-NGF promotes sperm motility and viability in humans [31, 32], bulls [33] and golden hamsters [34], and facilitates the acrosome reaction [34]. In rabbits, there are no studies about the effects of β-NGF in the seminal characteristics of ejaculated semen.

We hypothesize that specific substitutions within the amino acid sequence of β-NGF from rabbit may be relevant for its different physiological action and, consequently, the production and use of a homologous recombinant rabbit β-NGF can help to enlighten the specific role of this multi-functional protein as an OIF in this species. Moreover, the potential use of exogenous β-NGF in the seminal dose could be an assisted reproductive procedure to reduce the handling and costs at the insemination time in reflex ovulator species but effect in semen must be contrasted. In this context, the aims of the present study were: 1) to produce and purify recombinant β-NGF from the rabbit prostate amino acid sequence, verifying its biological activity in PC12 cells, 2) to compare and analyze β-NGF amino acid sequences from several representative species of induced and spontaneous ovulation, and 3) to assess recombinant β-NGF effects in sperm viability and motility in a dose-response study. All these aims were achieved; the protein was produced and purified and their effect in semen was successfully tested. Some relevant differences were found in the comparison of the β-NGF amino acid sequence among rabbit and the other species studied.

## Experimental procedures

### Production of recombinant rabbit β-NGF and functional assessment in PC12 cells

#### Animals, tissue extraction and processing

New Zealand White x California adult male rabbits were housed individually in flat-deck cages with a light program consisted of 16 h of light and 8 h of darkness, at 20 to 25 °C of temperature and a relative humidity of 60 to 75% maintained by a forced ventilation system. Each animal had free access to food and water. All the experimental procedures with animals were approved by the Animal Ethics Committee of the Polytechnic University of Madrid (UPM, Spain), and were in compliance with the Spanish guidelines for care and use of animals in research [35].

Animals (n=3) were euthanized and ventral media laparotomy was performed. Prostate complex was dissected [36]; the proprostate was discarded and the prostate part was recovered. Portions of 5 mm were collected in 1.5 mL tubes containing RNA later® Stabilization Solution (Ambion, Thermo Fisher Scientific, Washington, USA) to avoid RNA degradation. RNA later® was removed from tubes after being at 4° C overnight and samples were stored at −80° C.

#### Complementary DNA sequencing of rabbit prostate β-NGF

To the best of our knowledge, the cDNA sequence from rabbit prostate was not published, thus we proceeded to sequence it. Therefore, total RNA was isolated using TRIzol reagent (Life Technologies, Thermo Fisher Scientific, Washington, USA), and then mRNA was obtained with FastTrack® MAG mRNA Isolation Kit (Ambion, Thermo Fisher Scientific, Washington, USA) according to the protocol provided by the manufacturer. Afterwards, cDNA was synthesized using a mix of random hexamers (0.5 μg/μL) and oligo (dT) primers (0.1 μg/μL) (SuperScript™ First-Strand Synthesis System for RT-PCR, Life Technologies, Thermo Fisher Scientific, Washington, USA).

Firstly, to sequence the entire *NGF* gene, specific primers were designed (Table 1) for a highly conserved region of the gene among species. The design was performed taking into account that *NGF* presents alternative splicing; hence, amino acid and nucleotide sequences of different species were aligned (Clustal Omega Software and Serial Cloner 2.6 Software) to look for the conserved region. Polymerase chain reaction (PCR) was performed using 1 μl of cDNA as a template for *β-NGF* specific primers, using the Platinum® Taq DNA Polymerase kit (Invitrogen, Thermo Fisher Scientific, Washington, USA). Cycling conditions consisted of one first phase of 3 min for denaturation at 95 °C, followed by 40 cycles of 30 seconds at 95 °C, 30 s at 55 °C and 15 s at 72 °C, and a final phase of 5 min at 72 °C to allow elongation. Negative control without reverse transcriptase was performed in PCR, in order to discard genomic DNA contamination. A 2 % agarose gel was used to visualize the size of bands of the PCR products (10 μl per lane) by a scanner (Bio-Rad Laboratories, California, USA). The amplified products of 305 pb were purified from the agarose gel with SpeedTools PCR Clean-up kit (Biotools, B&M Labs, S.A., Madrid, Spain) and sequenced by Sanger method using the BigDyeTM Terminator v3.1 Cycle Sequencing Kit (Thermo Fisher Scientific, Washington, USA) in a 3730XL DNA Analyzer sequencer (Applied Biosystems, Thermo Fisher Scientific, Washington, USA).

**Table 1.**
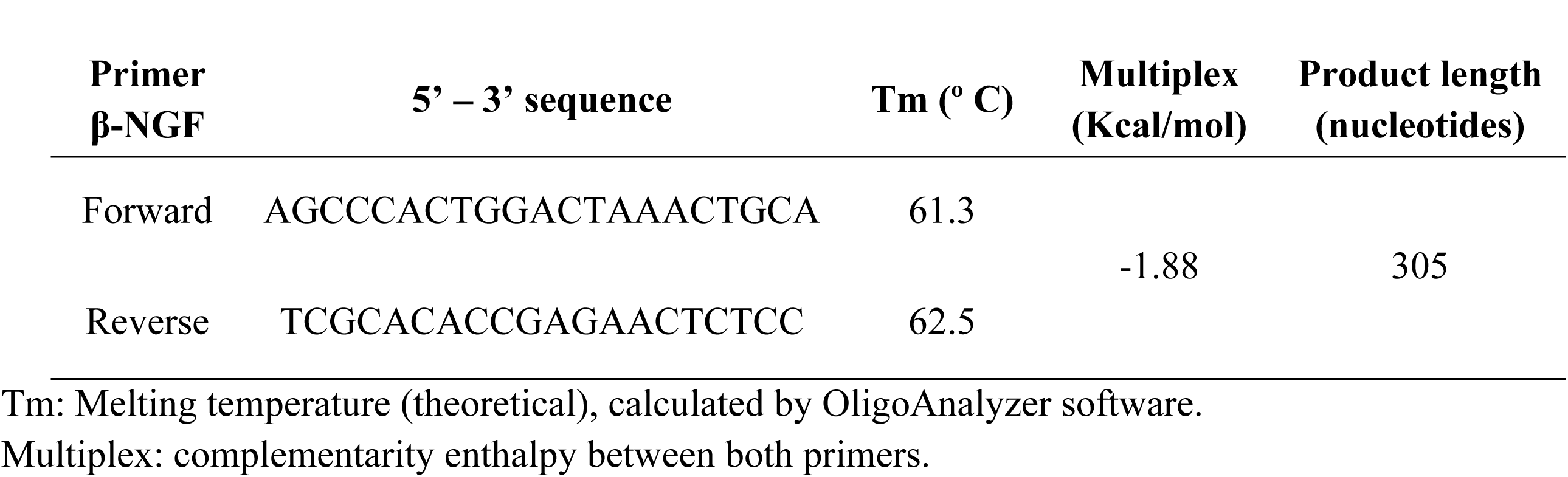
Specific primers for a conserved region of beta-NGF.

Once sequenced the conserved region of *NGF*, the Rapid Amplification of cDNA Ends (RACE) procedure was used to obtain the 5’ and 3’ sequences, based on Frohman et al. [37] using SMARTer® RACE 5’/3’ kit (Clontech Laboratories, California, USA). Inner and outer primers were designed following the kit protocol (Table 2), as well as the cycling conditions of PCRs, using a melting temperature of 68° C.

**Table 2.**
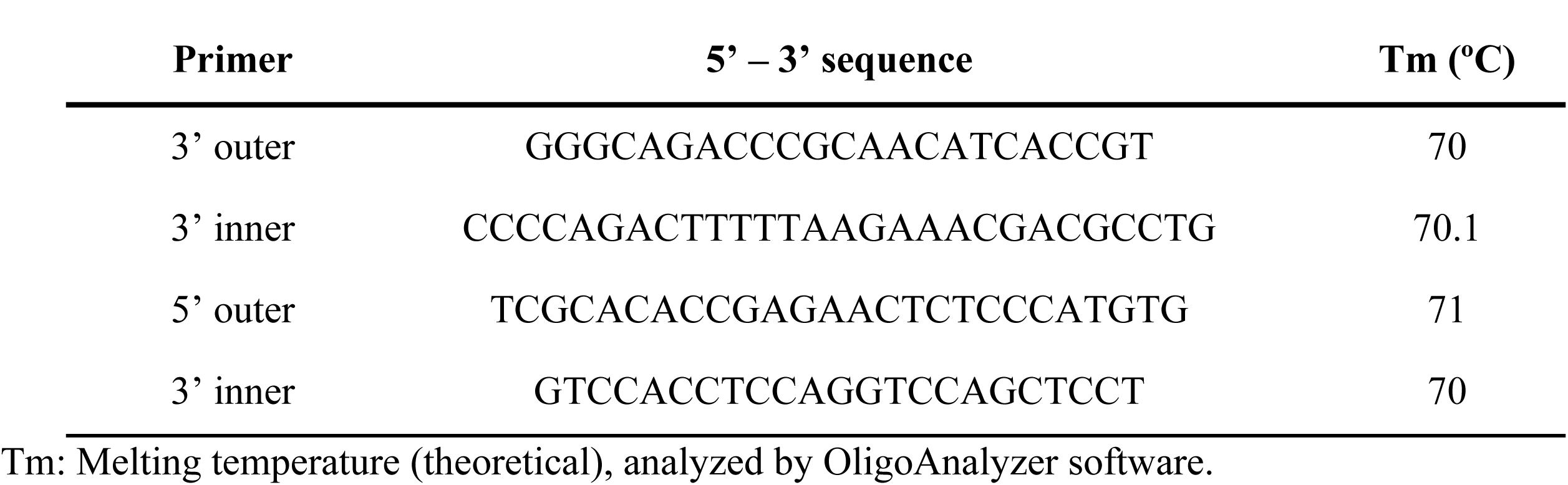
Specific primers designed for Rapid Amplification of cDNA Ends (RACE) procedure.

#### Plasmid production and transfection

Our rabbit prostate *NGF* sequence (KX528686) with a 7 × histidine tag in the C-terminal end was inserted in a pD2539-CEF plasmid with kanamycin and puromycin resistance (DNA 2.0, California, USA). Plasmid was produced in large quantities in bacterial cells (Stellar™ Competent Cells, ClonTech Laboratories, California, USA) using LB medium containing 25 μg/ml of kanamycin. They were incubated at 37° C by shaking during 24 h. Plasmid was extracted by Megaprep (PureLink® HiPure Plasmid Megaprep Kit, Thermo Fisher Scientific, Washington, USA) and then precipitated with isopropanol/ethanol. The resulting DNA was cut with the restriction enzyme Sal I to linearize the plasmid and then transfected with Lipofectamine® 2000 Reagent (Invitrogen, Thermo Fisher Scientific, Washington, USA) into Chinese Hamster Ovary (CHO) cells (ATCC, Virginia, USA).

#### CHO cells culture and protein purification

Transfected CHO cells were firstly cultivated in F12 medium supplemented with HEPES 25 mM (Gibco, Thermo Fisher Scientific, Washington, USA), 10% fetal bovine serum (FBS, Gibco® One Shot FBS, Thermo Fisher Scientific, Washington, USA), 125 μg/ml gentamicin (Invitrogen, Thermo Fisher Scientific, Washington, USA) and 5 μg/ml puromycin (Thermo Fisher Scientific, Washington, USA). Afterwards, 500,000 viable cells/ml were subcultured in a T-160 cc flask (Thermo Fisher Scientific, Washington, USA) in Serum Free Medium (CHO-S-SFM II, with hypoxanthine and thymidine, Thermo Fisher Scientific, Washington, USA) supplemented with 50 μg/ml gentamicin and 5 μg/ml puromycin for 4 days and shaken on a shaker platform. All cell cultures were incubated in a humidified incubator (NuAire, Minnesota, USA) at 37°C and 5% CO_2_.

Recombinant rabbit β-NGF (rrβ-NGF) purification from the culture medium was made by affinity chromatography, using columns containing Nickel (HisXL-Column High Density NICKEL, Agarose Bead Technology, Florida, USA) in order to select only those proteins with the histidine tag. Columns were equilibrated with 5 column bed volume of binding buffer (20 mM disodium phosphate, 500 mMNaCl, 10 mM imidazole at pH 7.5) and culture media was added, keeping in contact with the resin for 15 min. Then, after several washes of the column with binding buffer, the protein was eluted in 10 mL of elution buffer (20 mM Disodium phosphate, 500 mM NaCl, 500 mM Imidazole) and then dialyzed in HEPES 10 μM for PC12 bioassay, or in phosphate buffer saline 0.01M (PBS tablet, Sigma-Aldrich, Missouri, USA) for rabbit sperm bioassay in order to maintain semen viability. Dialysis was performed by shaking at 4° C, changing the medium 3 times with a minimum of 3 h dialyzing per time. Protein concentration was measured by the Bradford method [38].

#### Western Blot analysis

To verify the presence of rrβ-NGF, aliquots of dialyzed protein were subjected to TCA-acetone precipitation and then denatured in loading buffer (0.312 M Tris-HCl, 10% SDS, 25% 2-mercaptoethanol, 0.01% bromophenol blue, 50% glycerol) at 95°C. Afterwards, they were loaded on 12% SDS-PAGE gels and were run at 90 V for 2 h. One gel was stained using Coomassie (Sigma Aldrich, Missouri, USA) and the other one was used to transfer the protein to a nitrocellulose membrane (Ammersham^TM^ Hybond ECL Nitrocellulose Membrane, GE Healthcare Life Science, Barcelona, Spain) (80 mA per membrane for 80 min). Membrane was blocked during 1 h using Odyssey® Blocking Buffer (LI-COR Biosciences, Nebraska, USA), and then incubated at 4°C overnight with goat anti-NGF antibody (0.1 μg/mL) (N8773, Sigma-Aldrich) in blocking buffer with 0. 1% Tween 20. After several washes, membranes were incubated at room temperature (RT) for 1 h with secondary antibody (IRDye® 800CW Donkey anti-Goat IgG (H + L), LI-COR Biosciences, Nebraska, USA). After several washes, membranes were scanned with an Odyssey fluorescence scanner (LI-COR Bioscience, Nebraska, USA).

#### MALDI-TOF mass spectrometry analysis

For mass spectrometry analysis, SDS-PAGE and Coomassie staining were performed with the synthesized protein as described above, and gel bands of 13-15 kDa were manually excised from gels. The experimental procedure was executed as previously published [11]. For protein identification, the sequence of rabbit prostate NGF (KX528686) was searched using MASCOT v 2.3 (www.matrixscience.com) through the Global Protein Server v 3.6 from ABSCIEX.

#### PC12 cell culture

For PC12 cells culture, cells were thawed at 37°C, centrifuged at 1800 x *g* 5 min to remove DMSO from freezing and then plated in a T-75 cc flask. They were cultured in Dulbecco Modified Eagle Medium (DMEM, high glucose, HEPES) supplemented with 0.2 mM pyruvate, 10% horse serum (heat inactivated, New Zealand origin), 5% FBS and 50 μg/ml of gentamicin. All reagents were purchased from Thermo Fisher Scientific (Washington, USA). Medium was changed every 48 h, and cells were maintained in a humidified incubator at 37°C and 5% CO_2_. PC12 cells were seed at a density of 15,000 cells/500 μl in 24-well plates and grown for 24 h in a 37°C incubator. Culture media was added every 48 h with different concentrations of rrβ-NGF eluted in HEPES 10 μM: 0, 5, 10, 25, 50 and 100 ng/ml, respectively. Different concentrations were run in triplicates and each experiment was repeated 3 times.

#### MTT assay

To determine the possible cytotoxicity of rrβ-NGF treatment in PC12 cells, we assessed cellular viability at 48 h using 3-(4,5-dimethylthiazol-2-yl)-2,5-diphenyltetrazolium bromide (MTT, M6494, Thermo Fisher Scientific, Washington, USA) assay. After discarding the media of wells, 200 μl of 500 μg/ml MTT in Locke medium (140 mM NaCl, 4.5 mM KCl, 2.5 mM CaCl_2_, 1.2 mM KH_2_PO_4_, 1.2 mM MgSO_4_, 5.5 mM glucose, 10 mM HEPES) were added in each well and incubated for 2 h at 37°C. Then, 200 μl of solubilization buffer (0.1 M HCl, 1% Triton X-100 in isopropanol) were added and incubated for 1 h at RT to solubilize the formazan crystals. Sterile cell scrapers (Lab Clinics, Barcelona, Spain) were used to scratch the wells and the volume of each well was collected in tubes containing 0.9 mL of distilled water. The optical density (OD) of each sample was then measured at 560 nm using UltroSpec III spectrophotometer (Pharmacia LKB, GE Healthcare Life Science, Barcelona, Spain). The data were analyzed in terms of the percentage of cell viability, calculated by the equation: (OD treated cells / OD not treated cells) × 100.

#### Differentiation and neurite outgrowth assessment

In the moment of the bioassay, PC12 cells were plated in wells pre-coated with 7 μg/cm^2^ of collagen type IV (C6745, Sigma-Aldrich, Missouri, USA) at the same concentrations of rrβ-NGF eluted in HEPES 10 μM described before. In each concentration plate, 5 images of a minimum of 100 cells were taken with a light microscope (Leica F550, Wetzlar, Germany) equipped with phase contrast optics and a DCF400 camera (Leica). The morphological differentiation of these cells was assayed by determining the percentage of differentiated cells (loss of round shape), the percentage of cells with neurite elongations and by measuring the length of the longest neurite per cell at Day 8 with ImageJ software (https://imagej.nih.gov/ij/), according to Haas et al. [39]. Cellular elongations were considered as neurites when their length was one cell diameter at least [39].

#### Immunofluorescence against anti-β-III tubulin

The neuronal differentiation of PC12 supplemented with rrβ-NGF was evaluated by immunofluorescence against anti-β-III tubulin antibody, a microtubule element of the tubulin family found almost exclusively in neurons [40]. First, PC12 cells were cultured in 24-well plates in DMEM supplemented as described above, in covers treated with polylysine (Biochrom, Cambridge, UK). Twenty five ng/ml rrβ-NGF (the best concentration found in the previous analyses) were added 24 h after the beginning of the cell culture and changed every 48 h. At Day 8 of treatment, the medium was removed and cells were washed with 0.1M PBS (16 mM NaH_2_PO_4_2H_2_O, 84 mM Na_2_HPO_4_), fixed in 4% paraformaldehyde for 15 min at RT and treated with 0.1% Triton X-100 in PBS for 10 min at RT to permeabilize the cell membrane. After several washes, cells were blocked with 10% FBS in PBS for 45 min at RT to avoid non-specific binding and then, incubated with mAb against β-III tubulin (ab52623, Abcam, Cambridge, UK) at 1:50 dilution and 4°C overnight. Next, cells were washed with PBS and incubated with goat anti-rabbit IgG H&L (Alexa Fluor® 488) secondary antibody (ab150081, Abcam, Cambridge, UK) diluted 1:500 for 1h at RT. After washing in PBS, cells were incubated for 15 min at 4°C in PBS with 15 μg/ml Hoescht (B2261, Sigma-Aldrich, Missouri, USA). Covers were washed in PBS, rinsed in distilled water and mounted in a slide with mounting medium for immunofluorescence (VectaShield, VectaStain, Vector Laboratories, California, USA). The samples were observed by laser-scanning confocal microscopy (Leica TCS SP5, Wetzlar, Germany), using a 351/364 and 488 nm excitation lasers for visualize Hoescht and anti-β-III tubulin, respectively.

#### Trk receptor inhibition assay

To assure that rrβ-NGF was the responsible for PC12 cell differentiation into neuron-like cells, a specific inhibitor of the tyrosine protein kinase activity of the tyrosine kinase family (K-252a) was used in order to selectively block the effect of β-NGF in these cells. PC12 cells were incubated as previously described and K-252a (Sigma Aldrich, Missouri, USA) was added 24 h post-seeding at 100 nM and incubated for 2 h at 37°C. Then, three experimental groups were allocated: A) non-treated cells, B) cells treated with 25 ng/ml rrβ-NGF and, C) cells treated with 25 ng/ml rrβ-NGF and k252a, as mentioned before. The percentage of differentiated cells was assessed after 48 h of challenge with Trk inhibitor.

#### Study of the amino acid sequence of rabbit β-NGF

Full protein sequence of the rabbit prostate NGF (KX528686) was aligned with amino acid sequences of some species of induced ovulation (*Camelus dromedarius*, *Camelus bactrianus*, *Vicugna pacos*) and some species of spontaneous ovulation (*Rattus norvegicus*, *Mus musculus*, *Bos taurus*, *Homo sapiens*) using Clustal Omega Software. Glycosylation sites, disulfide bond sites, the signal peptide, the pro-peptide, the beta chain and the receptor binding sites were indicated in the output results of Clustal and were compared among species.

### In vitro assessment of sperm viability and motility with different concentrations of rrβ-NGF in semen samples

#### Animals, facilities and semen extraction

Six New Zealand White x California male adult rabbits (*Oryctolagus cuniculus*) held on the experimental farm at the Agrarian Production Department, Polytechnic University of Madrid (Spain), were used in this experiment. Semen was collected every week routinely in the farm by artificial vagina with a sexually receptive female.

#### Addition of rrβ-NGF to semen samples

After removing the gel fraction, mass motility of each sample was assessed and samples with the highest values were pooled to a final mean concentration of 383.4 ± 71.4 × 10^6^ sperm/ml and submitted to different doses of rrβ-NGF, depending on the experimental group. The doses were chosen according to β-NGF concentrations found in seminal plasma [10, 41, 42]. Hence the experimental groups were: 0, 2, 20 and 100 ng/ml of rrβ-NGF. PBS was added in 0 ng/ml group instead of rrβ-NGF (negative control group).

#### Sperm viability and motility analysis

Sperm viability and motility were assessed after 0, 1 and 2 h of rrβ-NGF challenge. Semen samples of each group at experimental times were collected for the assessment of sperm viability with nigrosin staining. Sperm motility was evaluated by Computer Assisted Semen Analysis (CASA), using the Motility module of the Sperm Class Analyzer (SCA®) version 5.2 (Microptic S.L., Barcelona, Spain). A minimum of 200 sperm cells per experimental group were recorded at each time. Then, parameters related to sperm motility were studied: static percentage (STAT, %), percentage of non-progressive sperm (NPMOT, %), velocity [curve-linear velocity (VCL, μm/s), straight-line velocity (VSL, μm/s), average path velocity (VAP, μm/s)] and percentages of linearity (LIN, %), straightness (STR, %) and wobble (WOB, %). This experiment was repeated 3 times.

### Statistical analysis

The data were analyzed using the SAS software version 9.0 (Statistical Analysis System Institute Inc, Cary, NC, USA). For the analyses of cell viability and neurite length of PC12 cells, a one-way ANOVA (GLM procedure in SAS) was used with β-NGF concentration (0, 5, 10, 25, 50 and 100 ng/ml) as fixed effect. For analyzing the differentiation and the neurite outgrowth of PC12 cells over time, we performed a two-way ANOVA with repeated measures (MIXED procedure in SAS), with β-NGF concentrations (0, 5, 10, 25, 50 and 100 ng/ml) and days of evaluation (2, 4, 6 and 8) as fixed effects, including also the interaction between these two fixed effects in the statistical model. To assess the effect of the protein concentration in the sperm, a two-way ANOVA was performed with β-NGF concentration (0, 2, 20 and 100 ng/ml) and time (0, 1 and 2 h) as fixed effects and the interaction between them was included in the statistical model too. All the variables are shown as mean ± s.e.m. and means were compared using Fisher test, considering significant differences when p-value < 0.05.

## Results

### Production of recombinant rabbit β-NGF and functionality in PC12 cells

The complete nucleotide sequence of *NGF* from rabbit prostate was sequenced from cDNA by RACE procedure and submitted to the GenBank database, with the reference number KX528686. rrβ-NGF was found to be expressed in the culture media of CHO cells transfected with the plasmid pD2539-CEF-rβ-NGF. A unique band of approx. 13-15 kDa was revealed by western blot. Furthermore, the corresponding protein band extracted from the SDS-PAGE gel once analyzed by fingerprint analysis combined with mass spectrometry (MALDI-TOF) showed a high score with the rabbit *β-NGF* gene sequence inserted into the plasmid (KX528686).

Cellular viability of PC12 cells was similar for all doses of rrβ-NGF challenged, except for the highest dose (100 ng/ml). This latter dose displayed a significant lower percentage of viability in comparison with rrβ-NGF doses of 5, 10 and 25 ng/ml (p<0.05) but similar to 50 ng/ml and 0 ng/ml doses that showed intermediate values (Fig 1A).

**Fig 1.**
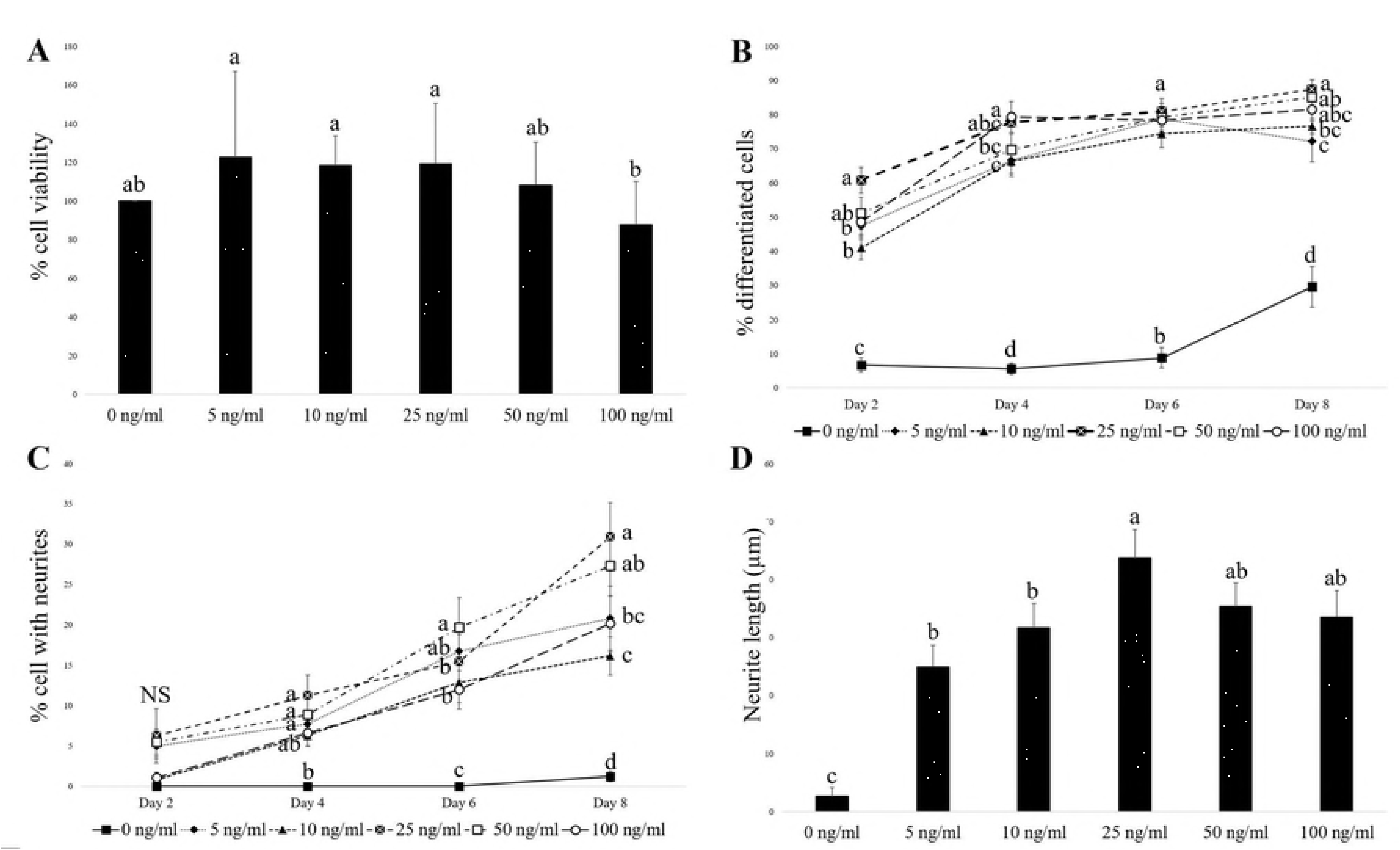
Dose-response study at different concentrations of rrβ-NGF (0, 5, 10, 25, 50 and 100 ng/ml) in PC12 cells. (A) Percentage of viability of PC12 cells after 48 h of culture at different rrβ-NGF concentrations. Different letters indicate significant differences between doses tested (p<0.05). (B) Percentage of differentiated PC12 cells over time (until Day 8) in culture with different rrβ-NGF concentrations. Different letters indicate significant differences in the same day between doses tested (p<0.05). (C) Percentage of PC12 cells bearing neurites over time (until Day 8) in culture with different rrβ-NGF concentrations. Different letters indicate significant differences in the same day between doses tested (p<0.05). (D) Neurite length (μm) of PC12 cells at Day 8 cultured with different rrβ-NGF concentrations. Different letters indicate significant differences between doses tested (p<0.05). All data are represented as mean ± SEM.

The percentage of PC12 cells differentiation was significantly higher in all rrβ-NGF concentrations groups than in the negative control group in all days evaluated (Fig 1B, p<0.05). At Day 2, rrβ-NGF 25 ng/ml triggered the highest percentage of differentiated cells, followed by the 50 ng/ml dose. At Day 4, cells treated with 25, 50 or 100 ng/ml presented the highest differentiation percentage, whereas the 10 ng/ml concentration group had the lowest one but still higher than the negative control. At Day 6, all rrβ-NGF treated groups presented the same differentiation percentage. At the end of the experiment (Day 8), the highest rate of differentiated cells was again found in 25, 50 and 100 ng/ml rrβ-NGF groups.

The percentage of cells with at least one neurite was not significantly different at Day 2 (Fig 1C). However, at Day 4, groups with 5, 25 and 50 ng/ml rrβ-NGF showed higher percentage of cells with neurites than 0 ng/ml group, whereas 10 and 100 ng/ml groups showed intermediate values. At Day 6 the percentage of cells with neurites was higher in 50 ng/ml group than all others, and 5 ng/ml group presented intermediate values, followed by the rest of rrβ-NGF groups. At Day 8, the rrβ-NGF treated cells with 25 ng/ml had the highest rate of cells with neurites, followed by the 50 ng/ml treatment.

Moreover, all rrβ-NGF groups showed longer neurites at Day 8 than the negative group (0 ng/ml dose). The average length of neurites was significantly higher in the group with the 25 ng/ml concentration than in the groups with lower concentrations (5 and 10 ng/ml), while cells treated with 50 and 100 ng/ml had intermediated average lengths (Fig 1D).

PC12 cells treated with 25 ng/ml of rrβ-NGF were positive for immunofluorescence with β-III tubulin at Day 8 of treatment (Fig 2) showing high fluorescence in all cytoplasm of the soma and in neurites. Cells cultured without rrβ-NGF did not show any expression of β-III tubulin.

**Fig 2.**
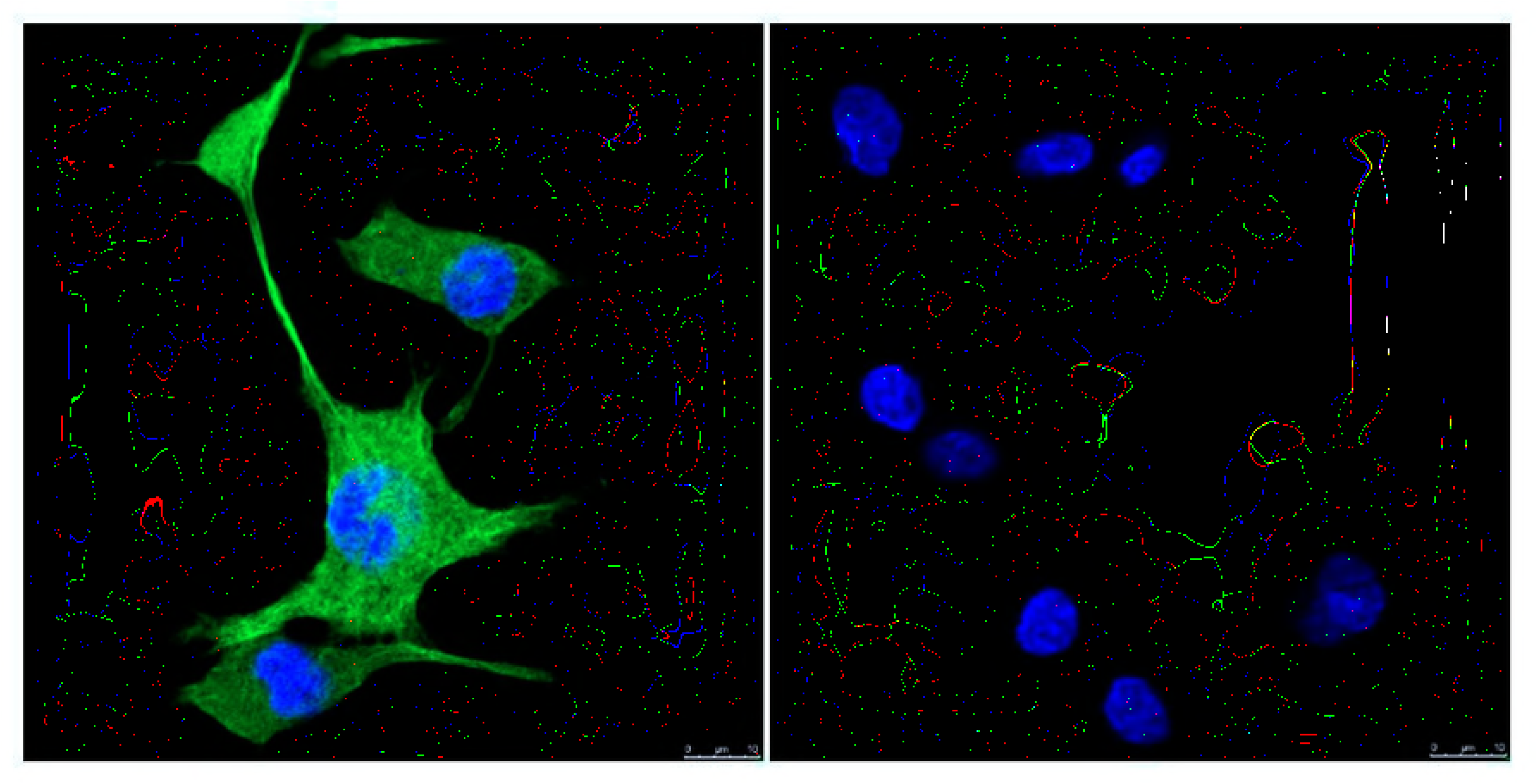
Immunofluorescence of PC12 cells treated with 25 ng/ml rrβ-NGF at Day 8 against β-III tubulin. Green signal represents the binding to β-III tubulin (soma and neurites) and blue signal is the nucleus stained with Hoescht 33342. Right panel is negative control (not incubated with primary antibody).

Finally, in the inhibition assay, PC12 cells did not show any differentiation or neurite growth after 48 h of culture in co-treatment with K-252a and rrβ-NGF (Fig 3). The percentage of cell differentiation in positive control group was significantly higher than in the negative control group (72.21±1.00 *vs*. 20.51±2.81%, respectively, p<0.05).

**Fig 3.**
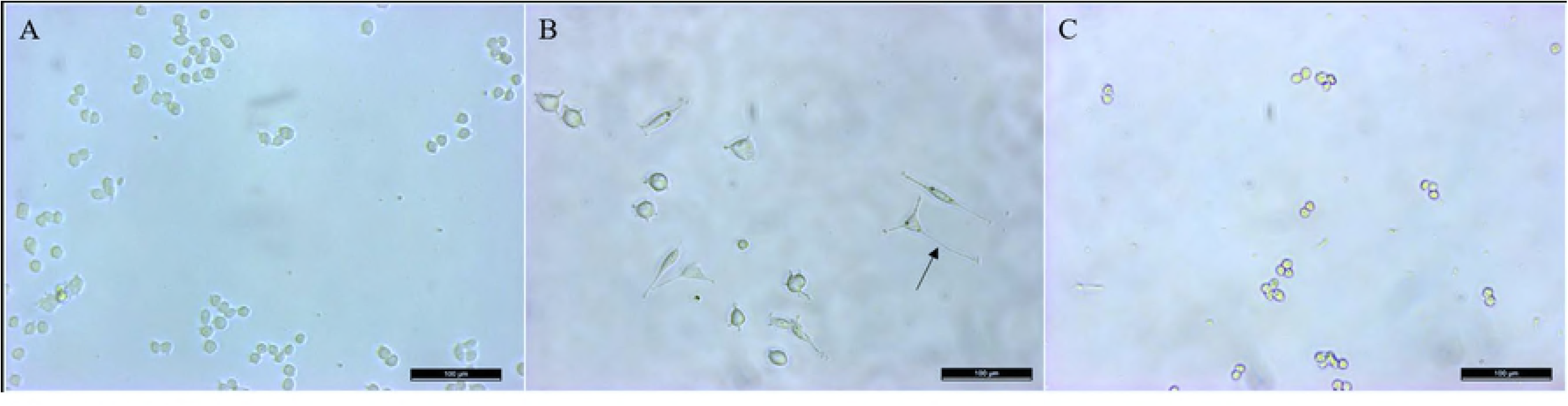
Inhibition of TrkA assay in PC12 cells. A: non-treated cells (negative control), B: cells treated with 25 ng/ml rrβ-NGF (positive control) and C: cells treated with k252a+25 ng/ml rrβ-NGF. Arrow: neurite. Scale bar: 100 μm.

### β-NGF protein sequence comparison among species

The amino acid alignment of rrβ-NGF with other species revealed that the signal peptide, the 3 glycosylation sites (Asn^69^, Asn^114^, Asn^166^), all the Cys that constitute the 3 disulfide bonds (Cys^136^ – Cys^201^, Cys^179^ – Cys^229^, Cys^189^ – Cys^231^) as well as Trp^142^ and Ile^152^, important for TrkA and p75 binding, were conserved in all species studied (spontaneous or not; Fig 4). In reflex-ovulation species, the N-terminal region of the beta chain of NGF, where β-NGF binds to its high-affinity receptor, presents the tandem Ala-Pro, whereas in spontaneous ovulators there is a corresponding Ser residue. Furthermore, specifically in rabbit, there is not a Ser after Ala-Pro, as in the rest of reflex-ovulation species. The majority of the amino acids related to the binding to p75 are conserved among species, except a sequence that is slightly different: KGNEVKVL in rabbit *versus* KGKEVMVL in the rest of studied species.

**Fig 4.**
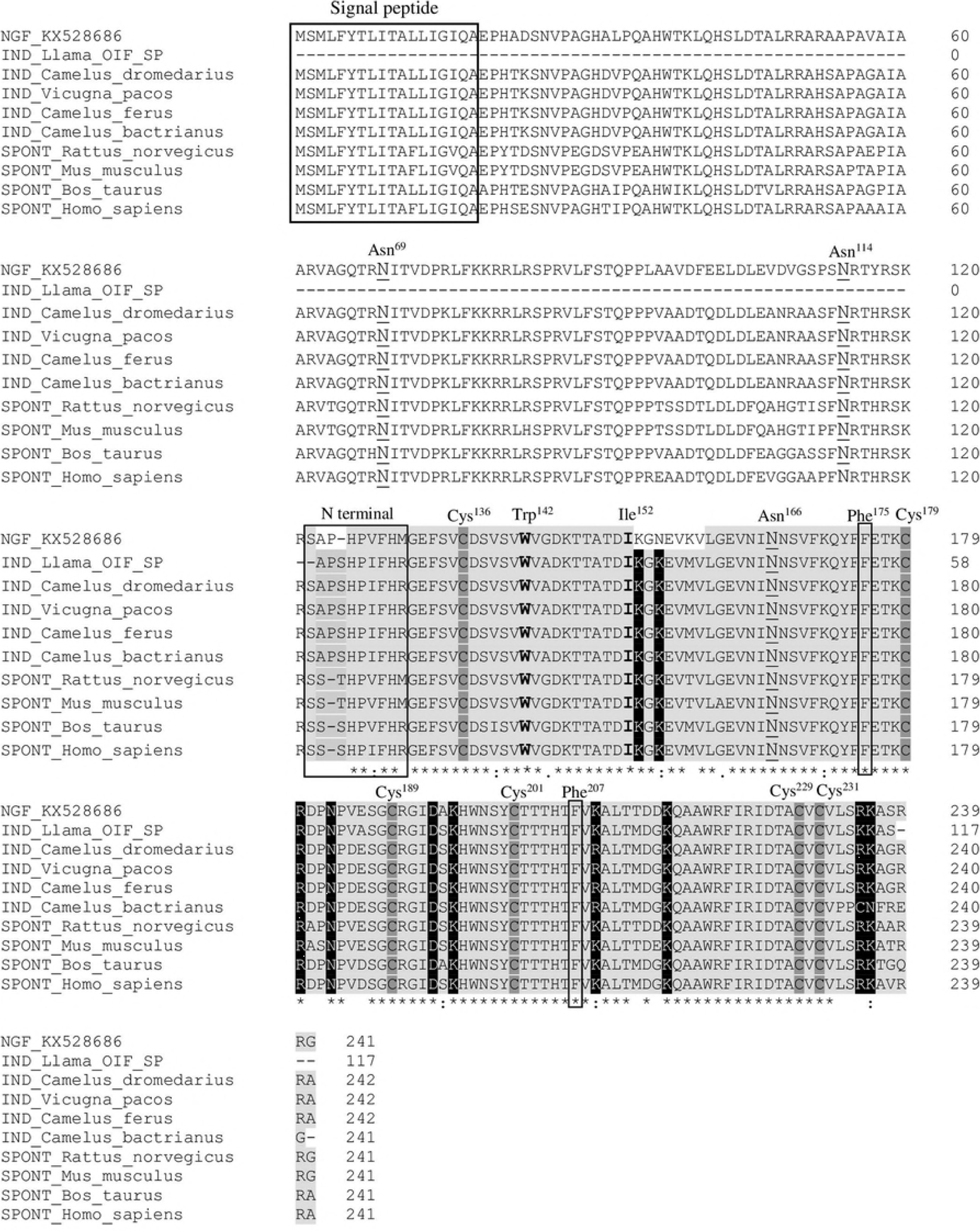
Amino acid sequence alignment of β-NGF from induced (IND, NGF_KX528686 from *Oryctolagus cuniculus*, KX528686; OIF from SP of *Lama lama*, 4EFV_B; *Camelus dromedarius*, XP_010979007.1; *Vicugna pacos*, XP_015102944.1; *Camelus bactrianus*, XP_010967135.1) or spontaneous (SPONT, *Rattus norvegicus*, NP_001263984.1; *Mus musculus*, NP_001106168.1; *Bos Taurus* NP_001092832.1, *Homo sapiens*, NP_002497.2) ovulation species. The signal peptide is indicated in a box. Underlying aa (N^69^, N^114^ and N^166^) show glycosylation sites of Pro-NGF. Beta chain of β-NGF is shaded in light grey. Cys involved in disulfide bonds are highlighted in dark grey (Cys^136^, Cys^179^, Cys^189^, Cys^201^, Cys^229^, Cys^231^). Amino acids that participate in both TrkA and p75 binding are in bold (Trp^142^ and Ile^152^). TrkA binding sites are indicated in black boxes (N-terminal, Phe^175^, Phe^207^). P75 binding sites are in white and highlighted in black. The differences observed in rabbit beta chain are in white color.

### Effect in rabbit sperm cells of rrβ-NGF addition in semen

Sperm viability (Fig 5A) and the percentage of static sperm (Fig 5B) were similar for all the concentrations of rrβ-NGF added to semen. CASA parameters related to the progression of the motion (NPMOT, Fig 5C; LIN, Fig 5G; STR, Fig 5H; WOB, Fig 5I) were affected by the highest concentration at 2 h, showing a decrease of progressive motion comparing to lower concentrations of rrβ-NGF. In addition, there were differences concerning to these parameters between 0 and 2 h in 20 and 100 ng/ml groups. However, the velocity parameters VCL (Fig 5D) and VAP (Fig 5F) presented higher rates in the 100 ng/ml group at 2h compared to groups with lower doses, and to 0h. In contrast, this group showed the lowest percentages for VSL.

**Fig 5.**
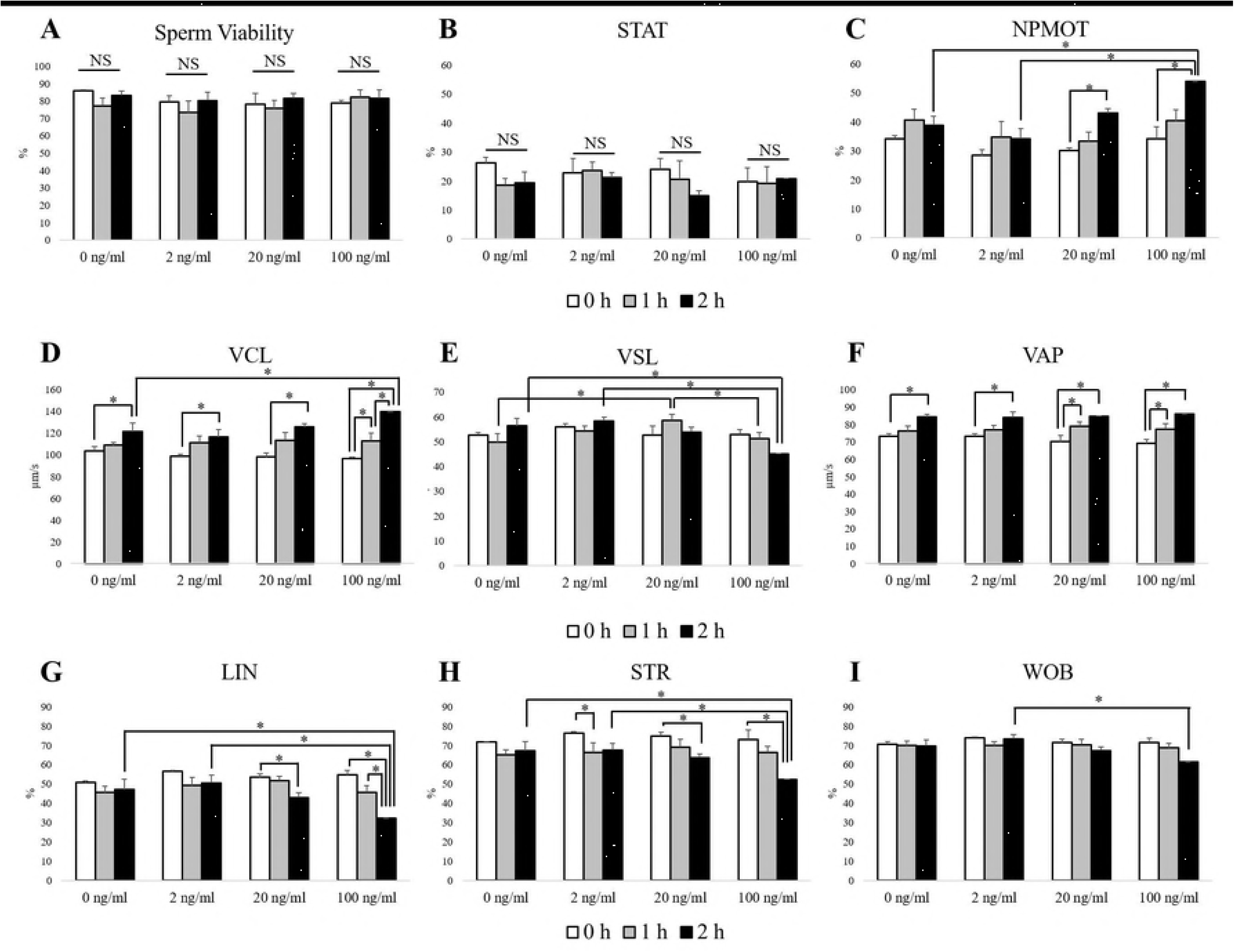
Sperm viability (A) and seminal motility parameters assessed by CASA (B-I) at 0, 1 and 2 h after addition of different doses of rrβ-NGF (0, 2, 20 and 100 ng/ml) to semen samples. Statistical differences are indicated by * (p<0.05). Data are represented as mean ± SEM.

## Discussion

In the present work, we have sequenced *NGF-mRNA* from rabbit prostate and produced and purified its corresponding recombinant rabbit β-NGF, which was able to differentiate PC12 cells into neuron-like cells. The comparison of the amino acid sequence of rabbit NGF with other induced and spontaneous-ovulator species has revealed species-specific differences, mainly in the receptor binding sites. Furthermore, we assessed the effects of the addition of this neurotrophin to rabbit sperm with the aim of verify the possibility of its addition in the seminal dose to improve the breeding systems in reflex ovulator species.

One of the challenges in the production of the recombinant proteins is the post-translational modifications, which can be achieved using the intracellular engineering of mammalian cells, such as CHO cells [3]. Thus, the transfection of the plasmid containing rabbit β-NGF in these cells resulted in the successful production of the native protein with an appropriate molecular weight (confirmed by western blot) and amino acid sequence (verified by MALDI-TOF), and the proper biological activity (cellular differentiation by neurite proliferation in PC12 cells). rrβ-NGF maintained PC12 viability regardless the concentration tested. However, 25 and 50 ng/ml treated cells presented the highest percentages of cell differentiation and neurite/cell during all the study, showing also a higher neurite growth as previously described [14]. In contrast, Gunning et al. [43] noticed that higher concentrations of mouse β-NGF than those used in the current study (150 ng/ml) can progressively increase the percentage of cells with neurites. Probably, the origin of β-NGF can affect PC12 cells differentiation due to the affinity to TrkA receptor [44].

Protein sequence of the rabbit β-NGF revealed some changes compared to other species in the binding sites to its receptors. Two consecutive residues Ala-Pro were found in the N-terminal region of rabbit β-NGF important for binding to TrkA, as in all species of induced ovulation. This association may indicate a different strength degree in the union to TrkA, since Pro has a special structure that may facilitate an angle of a greater torsion. Conversely, after this tandem of amino acids, a Ser residue is found in all of the species studied, except in rabbits. This missing residue may be relevant to create a more stable configuration through the torsion facilitated by the previous Pro. Furthermore, two amino acid residues which participate in the recognition of the low-affinity receptor p75 [45] presented also mutations in the rabbit sequence. Thus, Asn^155^ substitutes the conserved Lys, and Lys^158^ replaces also a conserved Met. The Lys residue has a positive charge and repels the protonated histidine within the binding site of p75. Its change to Asn residue, which has a neutral charge, may promote a closeness of β-NGF to its low-affinity receptor. We have recently reported the expression and localization of TrkA [46] and of p75 [20] in rabbit male genital tract evidencing that it probably has a role in rabbit reproduction. Despite the presence of these meaningful differences in amino acid sequence located at the binding domains to the receptors, some biological functions of the rabbit β-NGF appeared in parallel to other species. In llama females, rabbit seminal plasma produced ovulation at the same level as llama seminal plasma [16], thus the interaction with the receptor does not seem to be modified in other species. However, llama seminal plasma is not able to elicit the ovulation in rabbits [16], so it could indicate that these specific residues found in the rabbit amino acid sequence may explain in part some of the particular physiological characteristics in the rabbit ovulation process. In any case, it has to be taken into consideration the different sexual stimulation to trigger ovulation that occurs in both species and the high number of components of the seminal plasma. Further studies about mechanisms of rabbit β-NGF in the female reproductive tract are needed to elucidate its role.

β-NGF is present in the seminal plasma of several species and its high- and low-affinity receptors have been found in the head and tail of bovine [33] and human [47] sperm cells. Hence, the addition of β-NGF in semen presumably induces an effect in sperm. In the present study, the viability and the percentage of static sperm were unaffected by rrβ-NGF addition. In contrast, the motion pattern of the sperm was influenced in a dose- and time-dependent manner. The highest dose (100 ng/ml) and the longest time (2 h) in rrβ-NGF addition study in semen provoked a reduction of sperm progressivity. The data found in other species are hardly comparable to this study since the β-NGF concentrations and times of image captures were not equivalent. Nevertheless, in golden hamster, motility parameters at the time of the supplementation with 100 ng/ml of β-NGF did not seem to be affected compared with sperm without β-NGF [34]. In humans, the motility pattern and velocity appear to be improved with β-NGF doses of 1 and 10 μM and with 1 h of incubation [31, 48]. However, it is remarkable that rrβ-NGF had not negative effects in the moment of the addition and maintains the sperm viability over time. These findings could be interesting for the use of this neurotrophin in the seminal doses for experimental studies of ovulation or female fertility.

In conclusion, the differentiation of PC12 cells together with the appearance of β-III tubulin and the absence of neurite growth in the presence of TrkA inhibitor confirm that this novel recombinant rabbit β-NGF produced in CHO cells is a functional protein. This protein has some unique amino acid residues in the binding sites of the receptors, which may help to understand some of the particularities in the reproductive physiology of rabbit. In addition, the exogenous addition of rrβ-NGF to ejaculated rabbit sperm maintained viability and progressive motility of spermatozoa. Therefore, this new recombinant protein could be potentially used by the intravaginal via to elicit ovulation in rabbits and maybe in other reflex ovulator females.

## Author contributions

### Conceptualization

JM Bautista, RM Garcia-Garcia, PG Rebollar, PL Lorenzo.

### Formal analysis

A Sanchez-Rodriguez, RM Garcia-Garcia, PG Rebollar.

### Funding Acquisition

RM Garcia-Garcia, PG Rebollar, PL Lorenzo, M Arias-Alvarez.

### Investigation

A Sanchez-Rodriguez, P Abad, M Arias-Alvarez, RM Garcia-Garcia.

### Methodology

A Sanchez-Rodriguez, P Abad, JM Bautista.

### Supervision

RM Garcia-Garcia, PG Rebollar, PL Lorenzo, JM Bautista.

### Writing – Original draft preparation

A Sanchez-Rodriguez, RM Garcia-Garcia

### Writing – Review and Editing

All the authors

## Acknowledgements

We would like to thank Dr. P. Serranillos and Dr. Juan Tamargo (Dept. Pharmacology, Pharmacognosy and Experimental Pharmacology, UCM) for kindly providing PC12 cells and CHO, respectively. Also, we gratefully acknowledge B. Muñoz Velasco for its technical assistant in the rabbit farm, B. Aguado Zorrilla for helping in image analysis and L. Gutierrez from Proteomic Service of UCM for her invaluable help in this work.

